# *Rora* regulates activated T helper cells during inflammation

**DOI:** 10.1101/709998

**Authors:** Liora Haim-Vilmovsky, Johan Henriksson, Jennifer A Walker, Zhichao Miao, Eviatar Natan, Gozde Kar, Simon Clare, Jillian L Barlow, Evelina Charidemou, Lira Mamanova, Xi Chen, Valentina Proserpio, Jhuma Pramanik, Steven Woodhouse, Anna V Protasio, Mirjana Efremova, Julian L. Griffin, Matt Berriman, Gordon Dougan, Jasmin Fisher, John C Marioni, Andrew NJ McKenzie, Sarah A Teichmann

## Abstract

The transcription factor *Rora* has been shown to be important for the development of ILC2 and the regulation of ILC3, macrophages and Treg cells. Here we investigate the role of *Rora* across CD4+ T cells, both *in vitro* as well as in the context of several *in vivo* type 2 infection models. We dissect the function of *Rora* using overexpression and a CD4-conditional *Rora-*knockout mouse, as well as a RORA-reporter mouse. We establish the importance of *Rora* in CD4+ T cells for controlling lung inflammation induced by *Nippostrongylus brasiliensis* infection, and have measured the effect on downstream genes using RNA-seq. Using a systematic stimulation screen of CD4+ T cells, coupled with RNA-seq, we identify upstream regulators of *Rora*, most importantly IL-33 and CCL7. Our data suggest that *Rora* is a negative regulator of the immune system, possibly through several downstream pathways, and is under control of the local microenvironment.

## Introduction

Type 2 response is driven by a broad range of stimuli including helminths, allergens, specific bacterial and viral infections, and endogenous host molecules (*1*). This reaction is characterized by a cellular repertoire which include the innate lymphoid cells type 2 (ILC2) (*2, 3*) and adaptive T helper 2 (Th2) which are known to express transcription factor GATA-binding protein 3 (GATA-3). Those cells secrete distinct cytokines including interleukin-4 (IL-4), IL-5, and IL-13, driving eosinophil recruitment and immunoglobulin production.

Retinoic acid receptor-related orphan receptor alpha (*Rora*), is a nuclear receptor that functions as a ligand-dependent transcription factor (*4–6*), and is known to regulate functions in immunity in addition to other roles in development, circadian rhythm and metabolism (*7*). *Rora* was found to play a critical role in the development of ILC2 (*8, 9*) and for cytokine production in ILC3 (*10*). In the liver, it controls inflammation by promoting the macrophage M2 polarization (*11*). Together with *Rorc*, it influences the development of Th17 (*12*). Recently it was shown to have an important role in skin Tregs, where it is required for proper regulation of the immune response in atopic dermatitis. In this study, the deletion of *Rora* in Tregs resulted in exaggerated type 2 allergic skin inflammation, increased expression of *Il-10, Ahr* and *Il-4*, while *Tnfrsf25* expression was reduced (*13*).

Given this previous literature, we hypothesised that *Rora* might be involved in regulating an inflammatory response *via* CD4+ T cells more widely. To investigate this we assessed the function of *Rora* in four different infection models involving, but not limited to, type 2 immunity (Figure 1).

**Figure 1:**
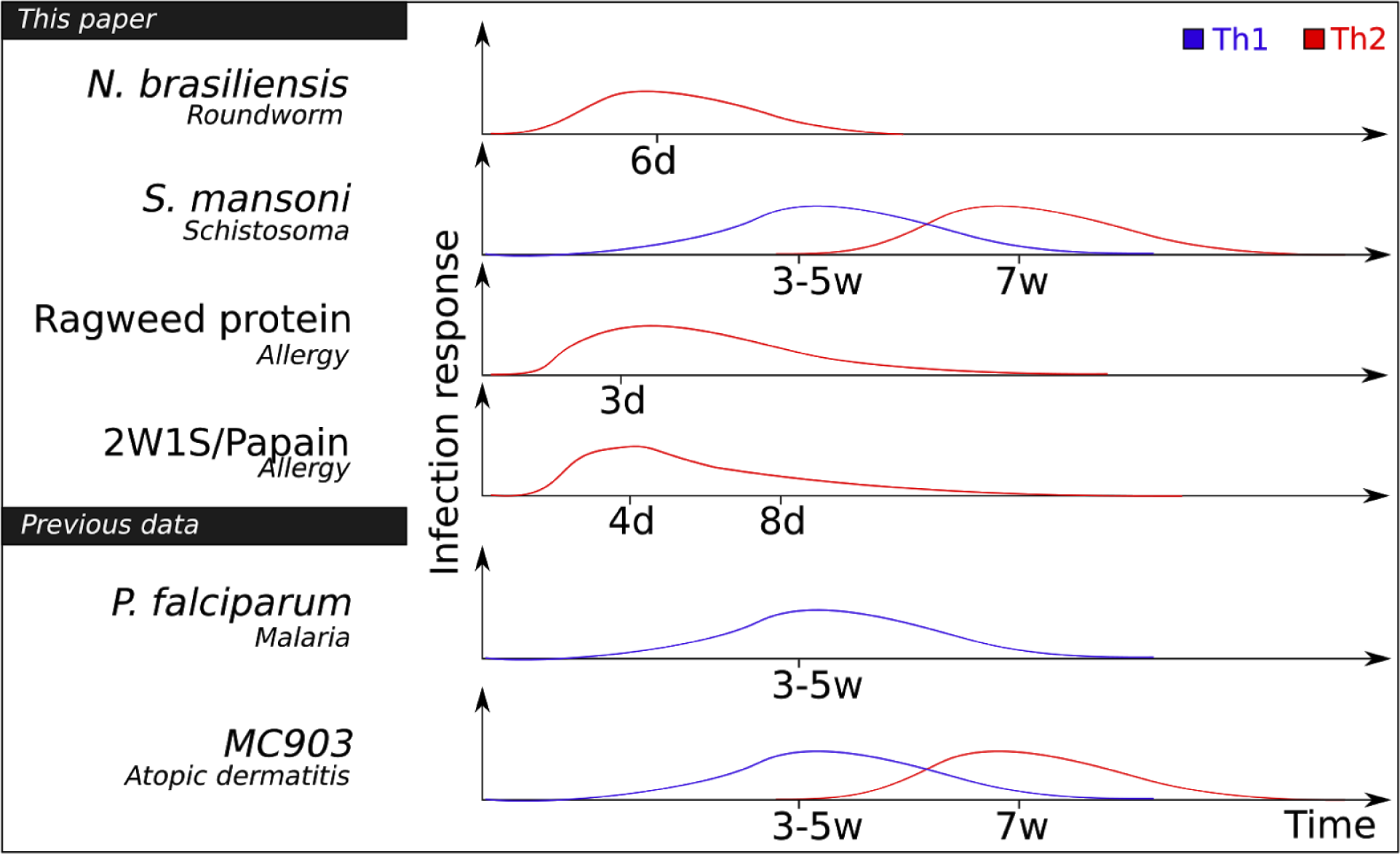
Overview of the experiments. The immune response to the infection models considered in this paper, over time, and which T helper cells are involved according to literature.

By employing bulk RNA sequencing (RNA-seq), single cell RNA-seq (scRNA-seq) and mass spectrometry (MS) analysis of T helper cells, we have investigated the role played by *Rora* in the context of *in vitro* cell culture and *in vivo* infection models. Our data suggest that *Rora* is expressed by activated CD4+ T cells, and its expression correlates with effector function, such as the expression of Th lineage-defining transcription factors and the production of cytokines.

## Results

### *Rora* is expressed in activated CD4+ T helper cells *in vivo*

To get an overview of *Rora* expression in CD4+ T helper cells during worm infection, mice were infected with *Nippostrongylus brasiliensis* (*N. brasiliensis*), a tissue migrating parasitic roundworm of rodents. *N. brasiliensis* initiates infection at the skin, the site of invasion, and then migrates via the circulatory system to the lungs. Through exploitation of host lung clearance, worms are transported to the intestine, from where they are expelled. In total, this infection is resolved within 7-10 days (Figure 1).

To get an overview of the infection at the molecular level, we performed single cell quantitative reverse transcription PCR (scRT-qPCR, Fluidigm Biomark). We prepared cells from the lungs, small intestine lamina propria (siLP), mediastinal lymph nodes (medLN) and mesenteric lymph nodes (mesLN) of mice, 3, 5 and 7 days after *N. brasiliensis* infection (Figure 2a). Tissues were also collected from uninfected mice as controls. Both CD3+CD4+SELL+ (naive) and CD3+CD4+SELL-(activated) cells were sorted. We examined 440 individual CD3+CD4+ T cells for the expression of 96 genes including housekeeping genes, different immune cell markers, T helper markers, cytokines, cytokine receptors, chemokine receptors, known and candidate regulators for Th cell differentiation (Figure 2b; Supplemental File S2). Consistent with a predominantly type-2 immune response, 70% of the CD3+CD4+SELL-cells expressed *Gata3*, whilst 26% expressed *Foxp3* and only 2% contained *Tbx21* or *Rorc*. Interestingly, in this population, we noted a significant overlap of *Gata3* and *Rora* (Fisher, p=4*10^−4^), and also of *Rora* and *Foxp3* (Fisher, *p*=2*10^−4^) (Figure 2b).

**Figure 2:**
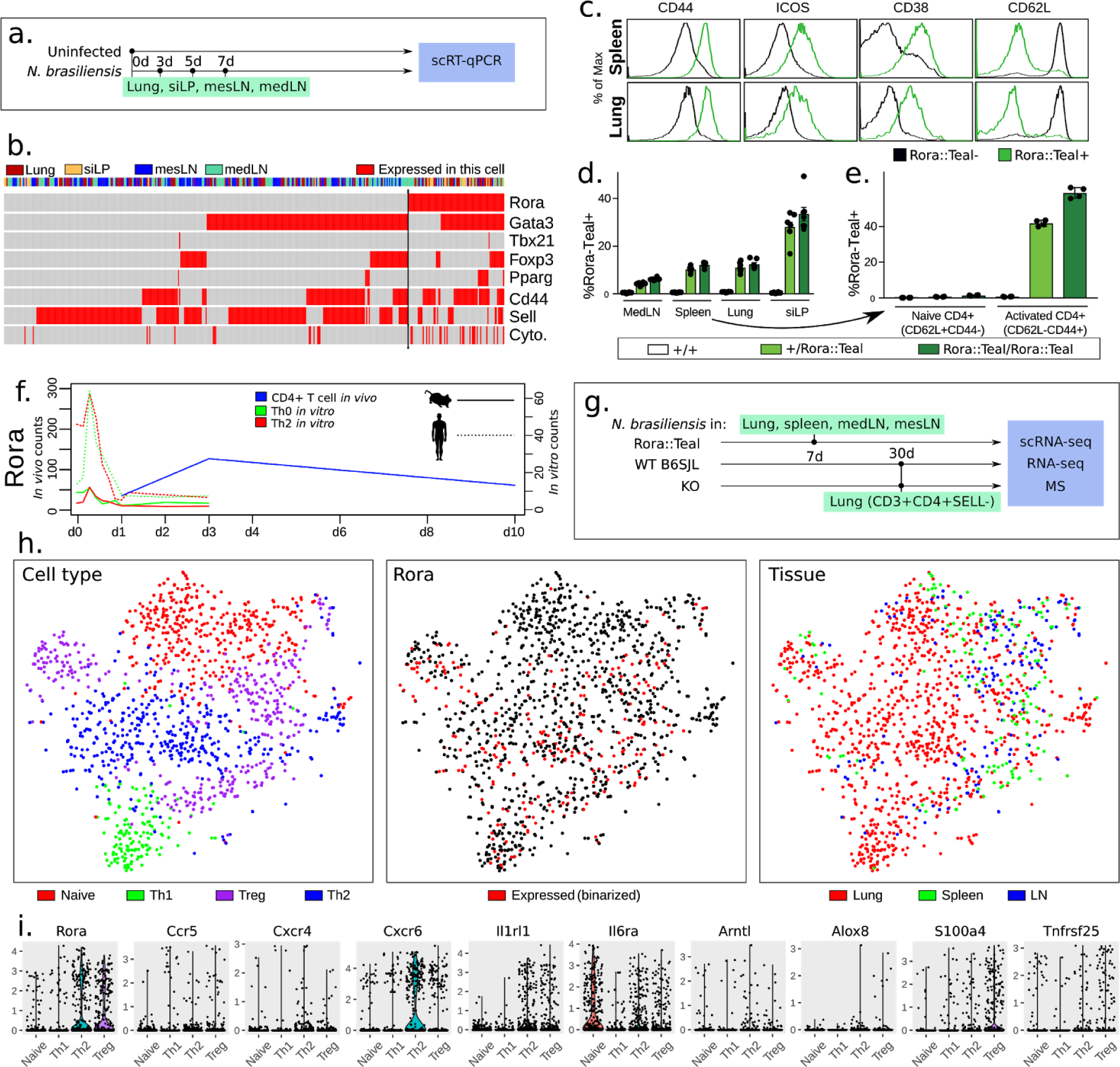
*Rora* is expressed in activated T cells *in vivo*, with mainly spatial variation. **(a)** Experimental design for single-cell RT-qPCR. **(b)** Gene expression according to RT-qPCR from CD4+ T cells during *N. brasiliensis* infection. *Gata3*^hi^ cells, which we assume are Th2, are also expressing *Rora*. The category “Cytokines” mean that any of *Ifng, Il4, Il13, Il5, Il6, Il10* or *Il17a* are expressed. Top row denotes spatial location: Lung, gut (siLIP), mesenteric and mediastinal lymph nodes (n=4 mice). **(c)** Flow cytometric analysis of CD4+ T cells from the spleen and lung of Rora^+/teal^ mice (black histogram = Rora^teal^– cells, green histogram = Rora^teal^+ cells) (n=23). **(d)** Frequency of Rora^teal^ expression across tissues, with siLIP having the highest number (n=12). **(e)** Fraction of naive or activated cells expressing Rora^teal^. RORA is mainly expressed in activated cells *in vivo (n=12)*. **(f)** *N. brasiliensis Rora* expression over time, overlaid with previously published mouse and human *in vitro* time course data of *Rora*, showing no or low expression of *Rora* in activated *in vitro* T cells. This suggests that a *Rora*-inducing element is missing in the *in vitro* system. **(g)** Experimental design of scRNA-seq experiment, with **(h)** tSNE clustering of CD4+ T cells extracted from *N. brasiliensis* infected mice and measured by single-cell RNA-seq (1670 cells after filtering). We primarily identify Naive, Th1, Th2 and Treg cells, of which *Rora* is expressed in all non-naive cells. **(i)** Expression distribution of selected genes from the scRNA-seq data.

To investigate *Rora* activity in activated Th cells, we examined *Rora* expression using a *Rora*^*+/teal*^ reporter mouse, in which the gene encoding Teal fluorescent protein has been inserted at the 3’ end of the *Rora* coding sequence and expressed from a self-cleaving T2A site (*14*). Flow cytometric analysis of reporter gene expression indicated that *Rora* was expressed in a proportion of CD4+ T cells isolated from medLN, spleen, lung and siLP (Figure 2c). We noted that all *Rora* expressing cells possessed an activated phenotype, as evidenced by their expression of CD44, ICOS and CD38 and down-regulation of SELL (Figure 2d). Accordingly, *Rora*^+^ T cells were therefore significantly enriched within the activated T cell subset, comprising 40-60% of this population, but absent from the pool of naive cells (Figure 2e). This correlation of *Rora* expression with T cell activation is in contrast with RNA-seq analysis made during the first 72h of *in vitro* culture of Th0 and Th2 cells, as previously reported (Figure 2f) (*15*). There, in both mouse and human, *Rora* is expressed in naive cells, with a sudden increase around 2-4 hours of activation, before a gradual decrease in expression (overall slightly higher expression in Th2 over Th0). This agrees with another *in vitro* activation dataset where the *Rora* expression level is unchanged and low, in both Th17 and Th1 cells, following activation (*16*). To compare with the *in vivo* situation, we generated RNA-seq data from CD4+ T cells, isolated from the spleens of *N. brasiliensis* infected wild-type mice 1, 3 and 10 days after infection, and overlapped with the *in vitro* data (Figure 2f). Here *Rora* is seen to increase, reaching a high and fairly sustained level around day 3. This is in agreement with a previous single-cell analysis of Th1/Tfh cells during malaria infection where *Rora* expression is seen to increase over time in Th1 (*17*) (Figure S1a, S1b). It is also consistent with a study on atopic dermatitis (*10*), a study on Tregs (*18*) (Figure S1c), as well as on T cells in a melanoma model (*19*) (Figure S1d). Taken together, as *Rora* is consistently present *in vivo* but not *in vitro*, these 6 datasets suggest that *Rora* is induced not primarily by activation, but by an external cue from the micro-environment.

Building upon this observation, we examined our scRT-qPCR data across different organs (Figure 2b). Overall, the expression of *Rora* was relatively constant in CD3+CD4+ cells 3-7 days after infection (Figure 2f). However, in terms of tissue, it was expressed more frequently in the siLP and lungs, where 37% of cells expressed *Rora*, compared to 20% of the cells in lymph nodes. This pattern was confirmed using flow cytometry, where 10.9±1.9% of CD4+ cells in the lung were RORA+, 10.1±1.1% from the spleen, 27.9±5.8% from the small intestine lamina propria and only 4.0±0.6% from medLN (Figure 2d). This tissue distribution likely reflects the relative frequency of activated Th cells in each location.

*Rora* has so far been seen in activated cells, partly overlapping with primary markers of Th2 and Treg (*Gata3* and *Foxp3*). To fully confirm the presence of *Rora* among different T helper cell types, as defined by the full transcriptional program, we used scRNA-seq on lymphoid and non-lymphoid tissue, isolating CD3+CD4+SELL-cells from the lungs 30 days after infection (*N. brasiliensis* infected wild-type mice 1, 3 and 10 days after infection, and overlapped with the *in vitro* data (Figure 2f). As reference points we included CD3+CD4+Rora^teal^+ and CD3+CD4+Rora^teal^-cells from *N. brasiliensis* infected *Rora*^*+/teal*^ reporter mice. Lungs, spleens, medLN and mesLN were also taken from infected and uninfected *Rora*^*+/teal*^ mice, 7 days after infection. The cells were clustered and each cluster annotated using common markers (*Sell, Ifng, Gata3, Foxp3*) as Naive, Th1, Th2 and Treg cells (Figure 2h, all cells are shown in Figure S1e). Cells from the lymph nodes and spleen are mainly Naive or Treg, otherwise the cell origin and T cell fate appear fairly uncorrelated. The distribution of *Rora* expression level across the different CD4+ T cell types was investigated (Figure 2h-i). Again, we note that *Rora* is expressed among all activated T cells (including Th1).

To confirm that this distribution of *Rora* expression is not specific to the response to *N. brasiliensis* infection we compared it to another worm model, *Schistosoma mansoni* (*S. mansoni*). CD3+CD4+SELL-cells were collected 6 weeks after infection from spleens and mesLN (Figure S1e). Single cell RNA-seq was performed and analysis shows that *Rora* is expressed in 59% of the activated cells (expressed defined as greater than zero counts). Here again, the approximately 30% cells that expressed *Rora* also expressed other key regulators (*Gata3, Foxp3, Tbx21*).

To conclude, we have performed FACS, scRT-qPCR and scRNA-seq, and confirmed that *Rora* is generally expressed in a proportion of activated T cells during *N. brasiliensis* infection. *Rora* induction is likely to depend on a local environmental cue not present in the usual CD3/CD28 *in vitro* activation system. Thus an *in vivo* system is required to study the physiological role of *Rora*.

### *Rora* is expressed in activated T helper cells during immune responses

To further corroborate these findings, we also investigated a non-worm immunity model, Ragweed pollen (RWP), a common allergen which causes lung inflammation (*20*). The *Rora*^*+/teal*^ reporter mice were exposed to intranasal RWP or PBS (Phosphate Buffered Saline, a negative control administration), administered every 2-3 days for 2 or 6 weeks (Figure 3a). As anticipated, RWP exposure resulted in increased CD4+ T cell counts in the lung and mediastinal lymph node (medLN) after 2 weeks as compared to PBS controls, and this was maintained after 6 weeks of continuous RWP administration (Figure 3a). In lung, but not medLN, this increase in CD4+ T cell frequency was accompanied by an elevated proportion of T cells expressing *Rora*, increasing from 30% to 40% between 2 and 6 weeks of RWP administration (Figure 3b and Figure S2). Consistent with our findings in untreated mice, RORA+ T cells elicited by RWP stimulation had an activated phenotype, characterised by a CD44+SELL-surface profile (Figure S2). Notably, the RORA+ T cell subset was enriched for cells that had acquired effector function, identified by their expression of the cytokines IFNg, IL-13 and IL-17A, or the Treg-associated transcription factor, FoxP3 (Figure 3c). Here again we note that *Rora* expressing cells include diverse activated cell phenotypes, that is, Th1, Th2 and Th17.

**Figure 3:**
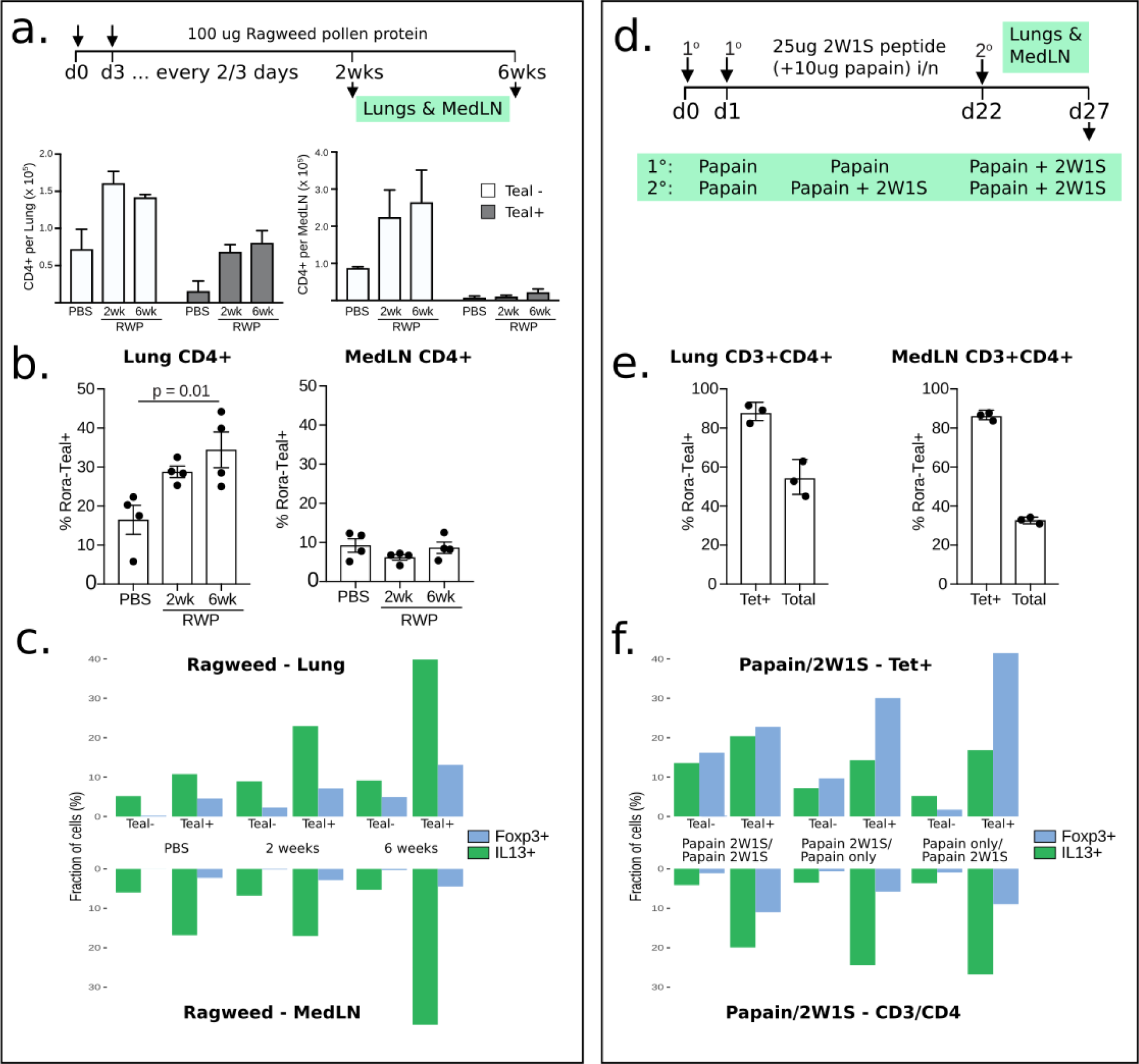
*Rora* is generally expressed in Th2 during type 2 immunity. **(a)** Flow cytometry analysis of CD4+ T cells from mice challenged with Ragweed pollen protein for 2 or 6 weeks, showing that *Rora* is expressed in lungs also during allergy. Here the number of Rora+/- cells are shown over time (n=12, one repeat). **(b)** RORA expression in CD4+ cells increases in the lungs but not in MedLN during the ragweed challenge (n=12, one repeat). **(c)** Flow cytometry analysis diagrams showing higher proportions of cytokines-expression and Tregs cells in Rora^teal^+ sorted cells *vs* the Rora^teal^-, after Ragweed pollen protein exposure (n=2). IL17 and IFNg were also measured (Figure S2b) **(d)** 2W1S/papain challenge experimental design. **(e)** Flow cytometry analysis when T cells are challenged twice with 2W1S/papain, again confirming the presence of *Rora* in tetramer+ cells. Note that we only performed this experiment once (n=3+3). **(f)** Flow cytometry analysis diagrams showing higher proportions of cytokines-expression and Tregs cells in Rora^teal^+ sorted cells *vs* the Rora^teal^-, after 21WS/Papain challenge (note that we only did this experiment once).

To understand the involvement of *Rora* in effector cells, we examined its expression in cytokine-expressing cells. From the *N. brasiliensis* model single-cell RT-qPCR, 83% of the cells that expressed either IL-4, IL-13 or IL-10, also expressed *Rora* (Figure 2b). The corresponding number in the *S. mansoni*-infected mice is 71% (Figure S1f, scRNA-seq)

To investigate whether *Rora* was expressed by Th cells responding to secondary immune challenge, we challenged *Rora*^*+/teal*^ and control mice with 2W1S peptide intranasally, in conjunction with the protease allergen papain, which promotes a type-2 immune response (Figure 3d). After secondary immunisation with 2W1S/papain 3 weeks after the initial stimulation, we were able to identify 2W1S-responsive CD4+ T cells by tetramer staining (*21*) and FACS analysis. Tetramer+ CD4+ T cells in both the lung and medLN were uniformly CD44+ and Rora^teal^+ (Figure 3e), indicating that recently activated Th cells express *Rora*. As with the *N. brasiliensis* infection, in the *S. mansoni* model we found significant overlap of *Foxp3* and *Rora* (Figure S8, Fisher, p=2*10^−4^). A higher fraction of FOXP3+ cells was also found in sorted RORA+ relative to RORA-cells, in the RWP model (Figure 3c). The gap is increased when the exposure to the pollen was longer. The same could also be seen after mice were exposed to 2W1S+papain. The RORA+ population contained 20% of the FOXP3+ cells, while the RORA-, only 4% of the cells.

Overall our data show that *Rora* is expressed in Th cells in worm immune models as well as in allergen type 2 immunity. We find *Rora* transcripts present in activated Th cells, not only in Tregs, but also in Th1 and Th2, as well as activated CD4+ T cells responding to a secondary immune challenge. *Rora* is also more correlated with cytokine-releasing cells, which suggests a function in effector cells.

### *Rora* KO affects inflammation severity

To isolate the *in vivo* physiological function of *Rora* in Th cells, we created a conditional knockout mouse line, where exon 4 of the *Rora* gene was deleted only in CD4+ cells (*CD4*^*cre*^*Rora*^*fl/fl*^, verified by RNA-seq, Figure S3a), resulting in a dysfunctional RORA protein. We noted no change in Th cell number when comparing *CD4*^*cre*^*Rora*^*fl/fl*^ to control (Figure S3b). We then infected *CD4*^*cre*^*Rora*^*fl/fl*^ mice with *N. brasiliensis* and examined lung inflammation after 30 days. An unbiased image analysis algorithm was developed to detect emphysema and score stained lung sections (Figure 4a). Generally, uninfected samples score the lowest for emphysema, as expected (Figure 4b,c). During infection, single-allele *Rora* KO (*CD4*^*cre*^*Rora*^*+/fl*^) had a higher score of emphysema, and double-allele KO (*CD4*^*cre*^*Rora*^*fl/fl*^) even higher score (linear model fit, *p*=0.018). To ensure that this is not a strain-specific defect we also compared infected WT mice with no-Cre KO mice (*Rora*^*fl/fl*^), as well as WT with Cre KO mice (*CD4*^*cre*^), and see no difference. It is possible that the emphysema is caused by eosinophil infiltration, in analogy with a previous model (*13*); we noted an increased but not statistically significant difference in eosinophil infiltration (Figure S3c).

**Figure 4:**
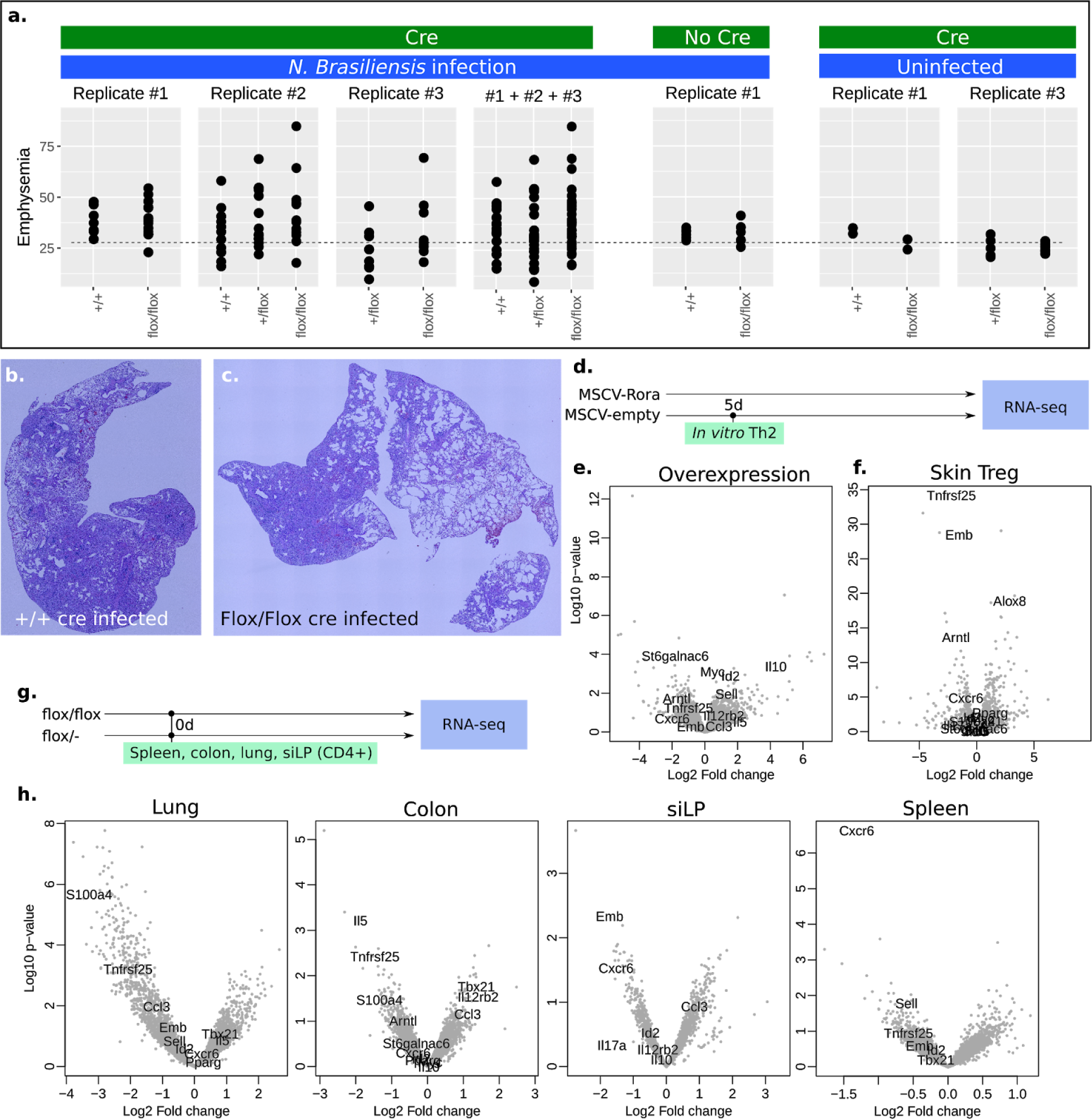
*Rora* KO affects lung inflammation severity. **(a)** Assessment of lung emphysema after *N*.*brasiliensis* infection by microscopy and automated image analysis. Controls are included for no-Cre (against the Rora-Flox) and no infection (from left to right, n=8, 17, 1, 14, 12, 9, 10, 29, 20, 39, 6, 6, 2, 2, 6, 6). **(b-c)** Least and most inflamed lung sections as detected by the algorithm, located in +/+ and Flox/Flox (control and KO). **(d)** Design of the *in vitro* overexpression experiment. **(e)** Differentially expressed genes with *Rora* overexpression. To increase readability, only some gene names are written out (full DE gene data is in Supplementary File S2). **(f)** Previous data, DE genes in Treg from skin (*13*) **(g)** Design of the *in vivo* KO experiment. **(h)** DE genes *in vivo*, activated CD4+ T cells (n=3+3), from lung, colon, siLP and spleen.

We conclude that *Rora* expression in CD4+ T cells modestly promotes lung inflammation associated with *N. brasiliensis* infection. This may reflect a role for *Rora* in regulating the activity of T helper cells, and may also result from the modulation of Treg activity, as has previously been reported for Rora+ Tregs in skin (*13*).

### *Rora* overexpression affects several key immune regulatory genes

To probe the role of *Rora* in T cells, we next proceeded to identify the molecular targets of *Rora* in Th2 cells by means of retroviral overexpression. Th2 cells were induced *in vitro*, infected the day after induction, and on day 5, gene expression was compared by RNA-seq (empty viral vector *vs* overexpression vector, Figure 4d,e). The *in vitro* differentially expressed (DE) genes were compared to the genes that are differentially expressed between skin Tregs from wt *vs CD4*^*cre*^*Rora*^*fl/fl*^ mice (Figure 4f). Consistent with a previous report (*13*), we see a strong effect on *tumor necrosis factor receptor superfamily member 25* (*Tnfrsf25)*, which has been shown to be required for Th2 effector function, *e*.*g*., allergic lung inflammation (*22*). The receptors *C-C chemokine receptor type 2* (*Ccr2*), *Ccr5*, and *Tnfrsf23* are in agreement, although with lower fold changes. We also find *Ninjurin 2* (*Ninj2*) which has recently been implicated in endothelial inflammation (*23*).

We also looked at a previous dataset of *Rora* small interfering RNA (siRNA)-treated human Th17 cells (*24*) and find that *Tnfrsf25* is DE (*p*=2.2*10^−4^). In another study of Tregs during steady-state in mouse (*25*), we find that *Tnfrsf25* is higher in colon Tregs than in lymphoid tissue Tregs (*p*=2*10^−2^). Thus four datasets, in different T helper cell types, confirm *Tnfrsf25* as a downstream gene.

Overall the two *in vitro* systems strengthen the claim of *Rora* as regulator of *Tnfrsf25*. However the expression of *Tnfrsf25* across Th2, Treg and Naive, but not Th1 (Figure 2i), shows that the function of *Rora* is not limited to Treg cells.

### *Rora* affects activation *in vivo*

To see if *Rora* has additional functions *in vivo*, we generated bulk RNA-seq from activated (CD62L-/CD44+) CD4+ T cells from control and *CD4*^*cre*^*Rora*^*fl/fl*^ mice from lung, spleen, siLP and colon (30 days after *N. brasiliensis* infection), and compared the DE genes between the different tissues, the overexpression DE genes, and the previous Treg *Rora* KO data. Volcano plots are shown in Figure 4 e-h. The full list of fold changes are in Supplemental File S2. From manual comparisons of these conditions, we have tried to find DE genes in common between the tissues, and if these in turn can explain the function of *Rora*. We generally do not see any DE cytokines of high statistical significance; we highlight some other types of genes here.

We find the transcription factor *Aryl hydrocarbon receptor nuclear translocator-like* (*Arntl*) to be differentially expressed in activated CD4+ T cells from siLP as well as in the overexpression analysis and in Tregs in skin (Figure 4f,h). *Arntl* is involved in the circadian responsiveness, and is expressed in CD4+ T cells (*26*). The *Arntl* promoter contains ROR elements (RORE), and was found to be regulated by *Rora* in mice (*27*). RORA promotes *Arntl* expression and thereby maintains the circadian rhythm. We find it to be expressed in all cell types (Figure 2i).

The *S100 calcium-binding protein A4* (*S100a4*) is another gene which is differentially expressed in activated T cells from lung, colon, and to lesser extent in Tregs in skin. In the single cell dataset, it is expressed mostly in Tregs but generally in Th1/Th2 (Figure 2h). It exists in both intra- and extracellular forms and can activate NF-κB (*28*). Using an antibody, *S100a4* has been shown to mediate T cell accumulation, and Th1/Th2 polarization, at tumour sites (*29*). Thus *S100a4* might act downstream of *Rora* to control T cell activation.

C-X-C chemokine receptor type 6 (*Cxcr6*) is differentially expressed in siLP, spleen and in Tregs in skin, and is consistently regulated in other tissues. *Cxcr6* is expressed in all Th cells but mostly in Th2 cells (Figure 2i). It might be involved in Th cell activation (*30, 31*).

The sialyltransferase *St6galnac3* is downregulated in colon and overexpression (Figure 4e,h). This enzyme transfers sialic acids to glycolipids and glycoproteins. Along with the previously DE gene *Alox8* (Figure 4f), we wondered if *Rora* may have any function on lipid metabolism. RORA has already been found to bind cholesterol (*32*) and is evolutionarily related to the nuclear receptor and lipid metabolism gene *Peroxisome proliferator-activated receptor gamma* (*Pparg*) (*33*). We performed lipid LC-MS on *in vivo CD4*^*cre*^*Rora*^*fl/fl*^ and WT cells from *N. brasiliensis* infected mice, as well as on *in vitro* WT naive and mature Th0/Th1/Th2 cells for reference (Figure S4). The *CD4*^*cre*^*Rora*^*fl/fl*^ presented small differences, and little overlap with the 85 lipids changed *in vitro* the first 6 days (cutoff *p*=0.05). However, all the top DE lipids are phosphocholines or phospoethanolamines, and some phosphocholines can induce a type 2 immune response (*34*). All the differentially expressed lipids are listed in Supplemental File S4.

In the spleen, *Sell* is downregulated in the KO, and opposite in the previous overexpression. This fits the idea of *Rora* driving activation and exit from the spleen, but why is it not DE in the other organs? We speculate that once a T cell has left the spleen or a lymph node, *Rora* might have already filled its purpose, and thus the effect on *Sell* will be smaller in any peripheral tissue.

We further noted the gene *Emb* (Embigin) in which is significantly DE in siLP, and in the same direction in overexpression, lung and spleen. It was also strongly DE in the previous skin Treg dataset. It has been shown to regulate cell motility in pancreatic cancer, and be controlled through the TGF-β pathway (*35*). It is possible that it could have a similar role in T cells.

Overall we see that *Rora* affects several genes *in vivo*, that are differentially expressed between different T helper cell types. Several of the downstreams genes may explain the effect of *Rora*, including for example *Arntl, Cxcr6, St6galnac3, Emb* and *Sell*. However, *Rora* abbrogration does not abolish T cells. It is possible that *Rora* acts as a helper factor, or tissue residence checkpoint, for T cell activation and migration.

### *Rora* expression is likely extrinsically regulated

A remaining question is which genes regulate *Rora*, and whether regulation is due to the T cell itself or from an external signal. Insight into when *Rora* is induced would provide understanding of its role. As previously seen, it is generally not expressed during *in vitro* activation, but is upregulated during *in vivo* activation. This discrepancy may be explained by an unknown signal from the environment that is present *in vivo* but not *in vitro*. To find candidate upregulators we screened for *in vitro* upregulated genes during 39 treatments, including cytokines and chemokines, with 15 unstimulated controls, in 3 biological replicates. In short, naive cells were taken from wild type mouse spleens, plated on CD3/CD28 coated plates, with addition of test cytokines/chemokines 24 hours later. 5 days after plating the cells were measured by bulk RNA-seq (Figure 5a).

**Figure 5:**
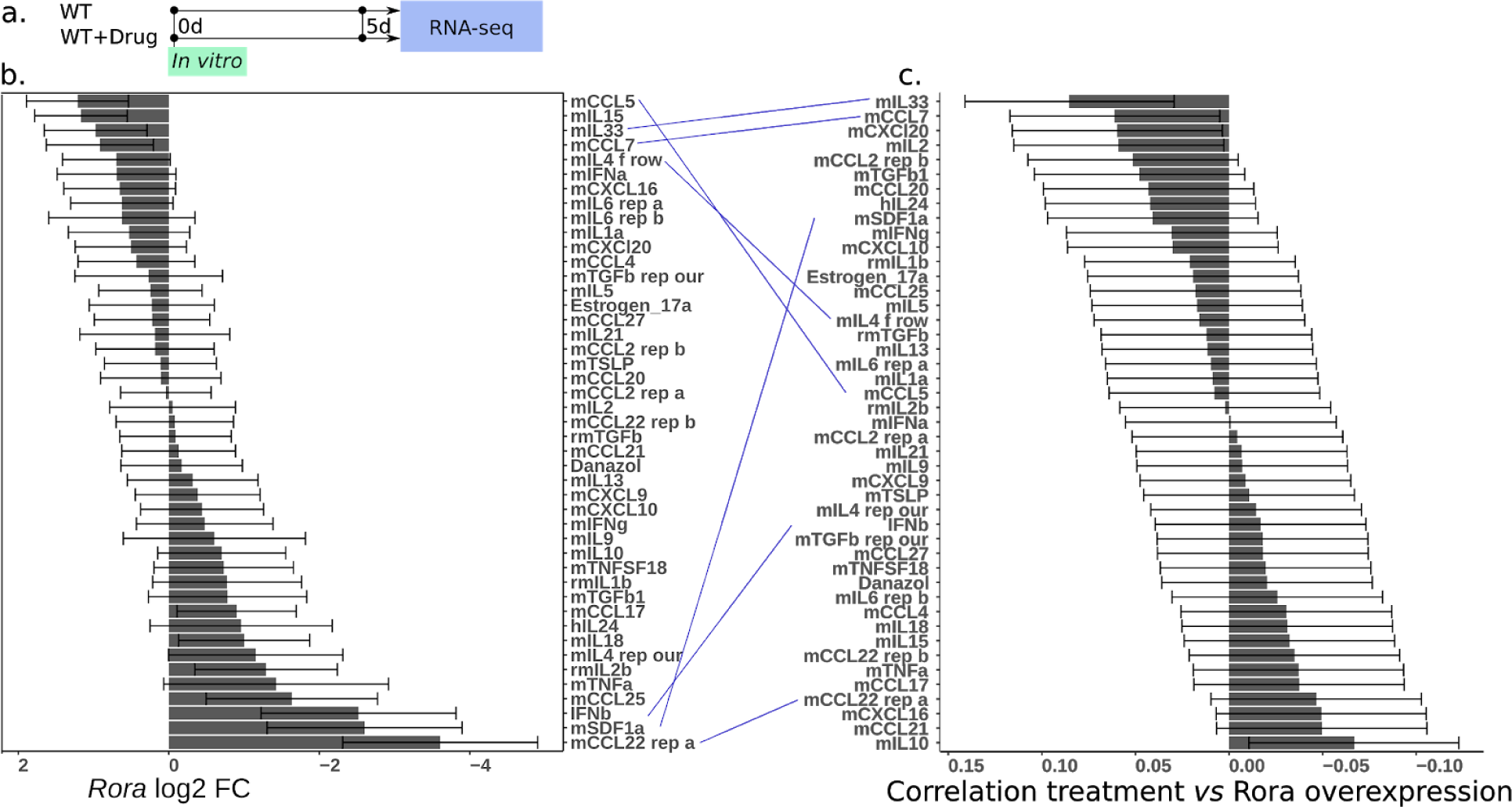
A screen of cytokine-treated T helper cells reveals genes upstream of *Rora*. The full list of treatments, and the effect on any gene of interest, is provided in Supplementary File S2. **(a)** Design of the *in vitro* experiment **(b)** Effect of all tested conditions on *Rora* expression. **(c)** Global correlation of gene fold change in each treatment, with fold change of gene expression during *Rora* overexpression. Lines have been drawn across panels b-c to highlight how similar a treatment is to directly perturbing *Rora*. Corresponding conditions are thus suggested to act through *Rora* as an early intermediate.

The stimulants and their effects are shown in Figure 5b (DE genes in Supplemental File S2). To validate our perturbation assay we looked at known marker genes. Of the genes having the strongest effect we find the expected genes: IL4 for *Gata3* (Th2), Transforming growth factor beta 1 (TGFb1) for *Foxp3* (Treg), IL6 and partially TGFb1 and IL21 for *Rorc* (Th17) (*36*) (Figure S5a,b,c). We then looked for the effect on *Rora* expression. The most upregulating stimulants are CCL5, IL15 and IL33.

CCL5 is chemotactic for T cells, eosinophils, and basophils. CCL5 signaling through CCR3 has been reported to regulate Th2 (IL4+CD4+) cellular responses to promote metastasis of luminal breast cancers (*37*). Interestingly also CCR5, another CCL5 receptor, was upregulated during overexpression, implicating a potential autocrine role for this factor. *In vitro*, CCR3 and CCR5 are expressed primarily in Th1, Th2 and Th17 (*16*).

IL15 is produced by non-lymphoid cells and important for the survival of several lymphoid subsets (*38*). IL15 KO mice are markedly lymphopenic due to decreased proliferation and decreasing homing to peripheral lymph nodes (*39*). Increase in IL15 thus fits our observation of *Rora* expression primarily in activated Th cells outside the lymph nodes. *Il15ra* is expressed in all murine CD4+ T cell subsets (*16*).

IL6 resulted in upregulation of *Rora* (statistical test significant if both batches of IL6 are pooled), consistent with previous IL6 *in vitro* culture data (*16*). IL6 has several roles, beyond generating Th17 cells *in vitro*, both pro- and anti-inflammatory (*40*). *Il6ra* is expressed in all CD4+ T cell subsets (*16*), though in our scRNA-seq data less in Th1 (Figure 2i).

IL33 also induces *Rora*, and is mainly associated with Th2 or ILC2 cells (*41*). This agrees with our *in vivo* scRNA-seq where *Rora* is slightly elevated in Th2 cells compared to other Th subtypes (Figure 2i). Our scRNA-seq shows that the corresponding receptor *Il1rl1* is expressed in all CD4+ T cell subsets but primarily in Th2 and Treg (Figure 2i).

*Ccl7* upregulates *Rora* and can do so through several receptors; these include for example CCR1, CCR2, CCR3, CCR5, and CCR10 (*42*). All of these receptors are present in CD4+ T cells to some extent, but *Ccr5* is somewhat higher expressed than the rest (*16*).

The most downregulating stimulants are C-C motif chemokine 22 (CCL22) and stromal cell-derived factor 1 (SDF1a). CCL22 is secreted by dendritic cells and macrophages, and is suggested to induce Treg cell infiltration into the pleural space in patients with malignant effusion (*43, 44*). In addition, its receptor CCR4 is required for CD4+ T cell migration to the skin (*45*). In bulk *in vitro* data it is present in all CD4+ T cell subsets, especially after activation (*16*).

SDF1a (also known as CXCL12), a ligand for the chemokine receptor CXCR4, promotes bone marrow homing for T cells. It has been shown that Naive T cells downregulate CXCR4 to avoid this (*46*). A further look at the Human Protein Atlas shows the tissues with expression of SDF1a to be particularly high in spleen, cervix/endometrium, and adipose tissue. In our study we compared with lung and gut, which have less *Cxcl12* expressed. This is consistent with our screening data, suggesting SDF1a may be a negative regulator of *Rora. Cxcr4* is expressed in all CD4+ T cell subsets at roughly equal level (*16*) according to bulk and our scRNA-seq data (Figure 2i).

To further validate if a treatment induces *Rora*, we also checked if a treatment induces the same downstream genes as in the *Rora* overexpression experiment. To do this we took genes from the *Rora* overexpression with a Log2 fold change > 2, and correlated the fold change with the corresponding fold change in a treatment (Figure 5c). Many of the most *Rora* influencing treatments also score similarly by this score. Different fold change cut-offs have been tested for reproducibility (data not shown). In particular, IL33 and CCL7 have a high level of agreement, followed by CCL22 (replicate a). Other treatments appear to have a less direct effect on *Rora*, for example IL6.

Based on our and previous data we conclude that *Rora* expression is mainly associated with an inflammatory state. Several cytokines may influence the expression of *Rora*. The effect on other genes can be studied using a similar screening approach, based on our supplementary data.

## Discussion

We have here investigated the impact of *Rora* on different subsets of CD4+ T cells, in several *in vivo* immune models. *Rora* is already known to be important for Th17 identity (*12*), and more recently it was discovered to regulate the function of Tregs in the skin (*13*). Here we show that *Rora* is actually expressed in all activated T helper cells during inflammation, including worm infections and allergy. We expand on the previous literature and suggest that *Rora* plays a wider role in the regulation of the immune response, in additional T helper cell types - Th1, Th2 and secondary activated cells as well. Its function is likely to also involve further key downstream genes than just the previously shown *Tnfrsf25*. Our model of the *Rora* regulatory network is depicted in Figure 6.

**Figure 6:**
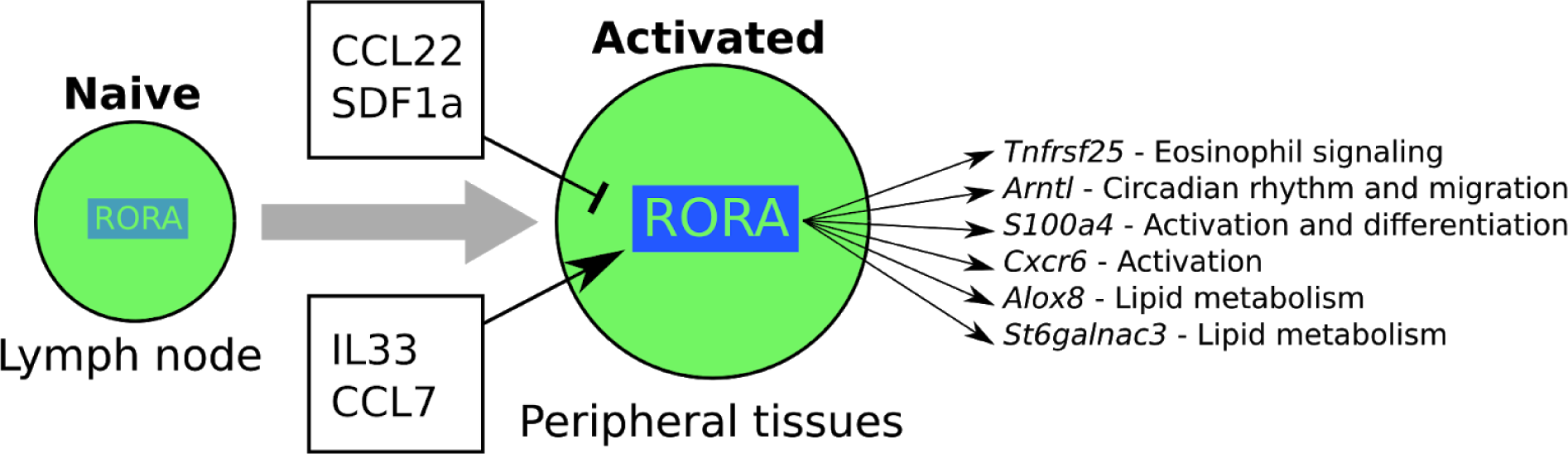
Summary model. *Rora* is expressed upon activation, and relies on additional stimuli from peripheral tissues for full induction of gene expression. Several cytokines have an impact on *Rora* expression level. *Rora* affects a diverse set of genes of importance for activated T cells, including *Tnfrsf25* (negative regulation of Eosinophils during infection), *Arntl* (circadian rhythm), *S100a4* (activation), *Alox8* and *Stgalnac3* (lipid metabolism).

*Rora* deletion did not alter the overall CD4+ T cell numbers, and therefore does not seem to be involved in T cell proliferation. This was also shown before in a study where Th2 cells were examined in RORα-deficient Staggerer mice (*47*). In this study we found that Th2 cells from the RORα-deficient mice were still able to differentiate and proliferate. However, we noticed higher inflammation in the lungs of *CD4*^*cre*^*Rora*^*fl/fl*^ mice, after *N. brasiliensis* infection, indicating a functional role in CD4+ T cells. A similar effect was shown in skin, when *Rora* was deleted in Tregs only (*13*). The effect was demonstrated to be through Treg *Tnfrsf25* signaling. *Tnfrsf25* has been shown to be required also for Th2 cell function in lung inflammation (*22*). Our analysis of the DE genes of activated Th cells, in the lung and colon of worm infected mice, as well as overexpression, and reanalysis of previously published data, confirms the importance of *Tnfrsf25* in Tregs, but that its origin may include also other activated T helper cell subsets.

A higher fraction of activated T helper cells (both Th and Tregs) contained *Rora*, when located in non-lymphoid tissues (lung and siLP) compared to lymph node tissues. The same pattern was shown previously for skin-located *versus* LN-located Tregs (*13*). The enrichment of Rora+ cells in peripheral tissues may reflect its correlation with T cell activation and effector function, or alternatively may relate to a role in influencing T cell migration. Our data is also unable to conclusively show if *Rora* is expressed before or after cytokines are expressed. Further research is required to see how *Rora* is involved in these regulatory programs.

One theory is that *Rora* has a complex interaction with circadian rhythm genes. T cells migrate in and out of the lymph node throughout the day (*48*), and *Rora* is already known to be involved in circadian rhythm. Our data strengthens this claim as the circadian gene *Arntl* is DE in both Tregs and other activated CD4+ T cells, in agreement with previous studies (*27, 49*). It is possible that *Rora* is regulated through external signalling. We investigated this with a small screen and found, for example, SDF1a as a negative regulator. This could be interpreted in the context of leukocyte migration during circadian rhythms, as blocking SDF1a disables the circadian migration of leukocytes (*48*).

Other theories can be brought forward based on downstream DE genes. If *Sell* is a direct downstream gene, it would explain most of the activation. However, other downstream genes such as *S100a4, Emb* and *Cxcr6* have also been linked to activation and cell migration. As *Rora* is a transcription factor which may have thousands of direct targets, the effect may be through more than one downstream gene, making it hard to pinpoint a single molecular mechanism.

The previous *Rora* KO sg mice display a range of phenotypes, *e*.*g*., reduced body weight gain and neuronal disorders. Assuming the gene regulatory network is preserved in other cell types, our work provides hypothetical mechanisms to explore further in future experiments. Since *Rora* controls PPARa (*50*), it may be a possible intermediate, with our data suggesting effects on phosphocholines. In this work we did not investigate the effect of potential *Rora* ligands but these could further contribute to a feedback loop. As for the neuronal disorders, knowing that the SDF1a-CXCR4 interaction is also important for neuronal guidance (*51*), SDF1a-mediated regulation of *Rora* may also explain neuronal impacts.

In conclusion, *Rora* has been found to be important for several immune cell types, in different contexts. It is involved in multiple pathways and appears to link cell migration with cell cycle, but also affects the outcome of inflammation through, but not limited to, *Tnfrsf25*. In its natural context, *Rora* is mainly associated with non-lymphoid tissue and activated T cells. *In vitro* culture suggests that this is due to environmental stimuli, possibly SDF1a. By generating a large systematic transcriptomic characterization of cytokine effects on T helper cells, we have found several cytokines that may explain be responsible for *Rora* induction in other tissues. With the large number of conditions tested in this study we are one step closer to understanding the multiple functions of this gene.

## Acknowledgements

We would like to thank Helen Jolin for help with mice, MRC Ares and Biomed staff, and Bee Ling Ng, Chris Hall, Jennie Graham, Maria Daly, Fan Zhang and Martyn Balmont for help with cell sorting. Ayesha Jinat provided technical assistance with RNA-seq.

## Author contributions

L.H.V designed the experiments, performed cell isolations, qPCR experiments and analysis, prepared cells for single cell RNA-seq experiments, bulk RNA-seq experiments, adoptive transfer and FACS experiments, did the C1 scRNA-seq experiments, and co-wrote the manuscript. J.W designed experiments, performed tissue and cell isolations, and most of the flow cytometry experiments and their analysis. J.H helped design some of the experiments, analyzed the data, performed the bulk RNA-seq experiments, analyzed some of the FACS data, did the overexpression and cytokine screen experiments, microscopy and image analysis, and co-wrote the manuscript. Z.M and G.K helped analyze the scRNA-seq data. E.N helped prepare tissues for qPCR experiments. S.W helped analyse scRT-qPCR data. J.L.B helped with the Ragweed model. M.E helped classify NKT cells. L.M helped with the single cell RNA-seq cell classification. S.C did the schistosoma worm infection. X.C and V.P helped generally with the experiments. A.V.P. harvested tissue from schistosoma-infected mice. G.D provided the *Schistosoma mansoni* infection model and advised on the study. J.F helped with the biomark data analysis. J.M helped with and advised on the analysis. A.C performed the MS and corresponding sample preparation. A.M provided most of the infection models, performed the histology scoring, and advised. S.A.T advised and conceived the project.

## Data and materials availability

The sequencing data has been deposited at ArrayExpress (E-MTAB-7694, E-MTAB-8000, E-MTAB-8001, E-MTAB-8003), and the image data at Zenodo (https://doi.org/10.5281/zenodo.2554078). The R code used for the analysis is available on GitHub (https://github.com/mahogny/liorarora).

## Supplementary files

S1. Biomark single cell RT-qPCR data, raw data and list of TaqMan probes

S2. RNA-seq count data and condition matrices

S3. List of antibodies

S4. LC-MS lipid data

S5. Supplemental Figures

## Methods

### Mice strains

#### Rora reporter mice

Rora T2A Teal reporter mice (*14*) were generated briefly as follows: A reporter cassette, encoding a short Gly-Ser-Gly linker peptide, FLAG epitope tag, T2A self-cleaving peptide and Teal fluorescent protein, followed by a loxP-flanked Neomycin cassette, was inserted (directly upstream of the) *Rora* stop codon. Successful targeting of JM8 ES cells was confirmed by Southern blot and the Neomycin cassette was removed from the resultant mice by inter-crossing with a Cre recombinase strain.

#### Rora KO mice

STOCK Tg(Cd4-cre)1Cwi/BfluJ mice were crossed with Rora^tm1a(EUCOMM)Wtst^ mice. F1 progenies were crossed again, and F2 were tested for having at least one allele of Cd4-cre, and two alleles of *Rora-lox*.

#### Ethics statement

The mice were maintained under specific pathogen-free conditions at the Wellcome Trust Genome Campus Research Support Facility or the Medical Research Council (MRC) Ares facility (Cambridge, UK). These animal facilities are approved by and registered with the UK Home Office. All procedures were in accordance with the Animals (Scientific Procedures) Act 1986. The protocols were approved by the Animal Welfare and Ethical Review Body of the Wellcome Trust Genome Campus.

### *O*verexpression of *Rora* and analysis

*Rora* was cloned into a retroviral vector (M6-mCherry-Cter) as follows. cDNA was made from RNA extracted from mouse spleens, following the protocol of SuperScript II reverse transcriptase (Thermo Fisher Scientific #18064014). Primers were designed against *Rora* isoform ENSMUST00000113624 (fwd: ATGTATTTTGTGATCGCAGCGATGAAAGCTCAAATTGAA, rev: TTACCCATCGATTTGCATGGCTGGCTCAAATT). After PCR and gel purification, another round of PCR was done to add compatible overhangs (fwd: CGATaagcttATGTATTTTGTGATCGCAGC, rev: GaatgcggccgcCCCATCGATTTGCATGG). The product and backbone were digested with HindIII-HF and NotI-HF (NEB), purified, and ligated using T4 ligase (NEB). Several clones were picked and sequence verified by sanger sequencing (M6-Rora2-mCherry). Two other constructs were also constructed analogously for ENSMUST00000034766 (M6-Rora1-mCherry and M6-Rora1-GFP).

The virus production, activation and transfection was done as previously, with the addition of IL4 (*15*). The transfection was done into 3 biological replicates (JaxJ, 8 weeks old), each with the 3 constructs and the no-insert M6 as control. No puromycin was used for selection. Cells were harvested on day 4, FACS sorted, and libraries prepared using the Sanger institute standard pipeline. Sequencing was done on a HiSeq 2500.

The differentially expressed genes were called by DESeq2 (*52*) (normalized counts ∼ treatment + mouse). Each of the constructs were analyzed. Of these, one had *Rora* as the most DE gene and was retained for further comparison (M6-Rora2-mCherry). Counts and condition matrices are provided in Supplementary File S2.

### Mice infection and cell isolation

#### *Nippostrongylus Brasiliensis* infection

C57BL/6 female mice were subcutaneously injected with 100ul (300 live third stage *N. Brasiliensis* larvae per dose), over two sites. MedLNs, mesLNs, spleens, lungs, or siLP were taken from infected mice, 3, 5, 7, 10 or 30 days after infection as well as from uninfected mice. At each time point, cells were isolated from medLNs and mesLNs, by smashing the tissue through 70μm cell strainers. Lungs were incubated in collagenase D (0.72mg/ml, Amersham, Bucks, UK) for 30min, smashed through 70μm cell strainers, and suspended in RBC lysis buffer (eBioscience Ltd). Spleens were smashed through 70μm cell strainers, and suspended in RBC lysis buffer. siLP cells were isolated from colonic tissue as previously described(*53*). In brief, siLP cells were released by digestion of the tissue with RPMI/4-(2-hydroxyethyl)-1-piperazineethanesulfonic acid (HEPES) supplemented with 60 μg/ml DNaseI (Sigma), and 400 ng/ml of Liberase (Roche Applied Science, Burgess Hill, UK). Isolated cells were stained with conjugated antibodies for sorting (Supplementary File S3). For RT-qPCR and bulk RNA-seq experiments, single cells were sorted into the lysis buffer and were stored in -80°C.

#### *Schistosoma mansoni* infection

The complete life cycle of the parasite *S. mansoni* is maintained at the Wellcome Sanger Institute. Cercariae, the mammalian infective stage, were harvested by exposure of infected *Biomphalaria glabrata* snails to light for two hours in aquarium-conditioned water. Female C57BL/6 mice were infected with 300-mix sex *S. mansoni* cercariae via IP as described elsewhere (*54*). Mice were checked twice a day for any sign of health deterioration. After 6 weeks post infection, mice were culled with an overdose of anaesthetic, adult worms were perfused from the mouse circulatory system as described previously (*55*), and mouse spleen and mesLNs were recovered surgically and placed in PBS for further processing. Cells were isolated from tissues and were stained for sorting. Activated T cells, CD3+CD4+CD44+CD62L-, were sorted into a C1 chip and processed for RNA-seq.

#### Lung immune challenge models

Mice were anaesthetised by isofluorane inhalation followed by intranasal administration of Ragweed pollen (100 μg per dose; Greer Laboratories, Lenoir, NC), papain (10 μg per dose; Sigma) or 2W1S peptide (10 μg per dose; Designer Bioscience), dissolved in 40 μl PBS at the indicated time points. The PE-conjugated I-A(b) 2W1S tetramer was obtained from the NIH Tetramer core.

#### Cell sorting

Cells were sorted with a BD Influx Cell Sorter or SY3200 Synergy cell sorter (iCyt) and analysed on Fortessa or LSRII BD cell analyzers using FlowJo. Surface and intracellular stainings were carried out according to the eBioscience protocols. A full list of the antibodies used is provided in Supplementary S4.

#### Histology

Lung lobes were fixed for 48 hours in 10% neutral-buffered formalin (Sigma), washed in PBS and transported to AML Laboratories (Florida) for embedding, sectioning and staining with haematoxylin and eosin.

#### Microscopy and image analysis

Images were captured using a VHX keyence microscope, using objective #2D1510064 at magnification “200”. The 2D stitching feature was used to capture the entirety of each lung section. Two sections were imaged from each mouse, with focus automatically calibrated for each section. All images were acquired in one seating, with the same settings for brightness.

Image analysis was done in R using the EBimage package (*56*). In brief, pixels corresponding to background, cytoplasm and nuclei were manually annotated in one image. A classifier was set up to separate pixels into the 3 classes based on manual inspection of the RGB space.

To estimate emphysema, the distribution of inner holes was calculated by first detecting non-background area. A binary dilation operation was applied to filter out 1-2 pixel holes caused by improper tissue detection. Segmentation was then performed to find disjunct regions. The area of each region was calculated.

From the histogram H of hole sizes, we estimate the unevenness (indicating emphysema, compensating for overall compression of slides due to fixation) as S=H_99%_/H_80%_. Other ranges were tested and gave a similar outcome. Veins, surrounding space, and other artifacts are filtered out above the 1% top percentile, while the lower percentile reflects tissue shrinkage. Alternatively, the score can be interpreted as a measure of hole size unevenness/variance. To test for emphysema in the *Rora* KO we fit a linear model S ∼ replicate + genotype, with genotype=1 for Cre and +/Flox, 2 for Cre and Flox/Flox, and 0 otherwise (*p*=0.018). A simpler t-test of +/+ *vs* Flox/Flox, not using any of the +/Flox samples, and ignoring replicate batch effects, gives *p*=0.17. This is not significant but in the direction of the more inclusive linear model.

#### *In vivo* bulk RNA-seq

RNA was extracted with SPRI beads, followed by Smart-seq2. Read counts were estimated with Kallisto (*57*). DE genes were estimated by DESeq2 (*52*) after counts had been rounded to the nearest integer count. Simple control *vs* treatment linear models were used. Counts and condition matrices are provided in Supplementary File S2.

### Quantitative single-cell gene expression analysis

#### Single cell RT-qPCR measurement and analysis

Single-cell gene expression analysis was performed using BioMark 96.96 Dynamic Array platform (Fluidigm, San Francisco, CA) and TaqMan Gene Expression Assays (Applied Biosystems, Carlsbad, CA). Single cells were sorted into 5μl of CellsDirect reaction mix and immediately stored in -80C. Control wells containing no cells were included. On thawing, a mix containing 2.5μL gene specific 0.2x TaqMan gene expression assays (Applied Biosystems), 1.2 μL CellsDirect RT/Taq mix, and 0.3 μL TE buffer were added to each well. RT–PCR pre-amplification cycling conditions were: 50°C, 15min; 95°C, 2min; 22x(95°C, 15s; 60°C, 4min). Samples were diluted 1:5 in TE buffer and 6% were mixed with TaqMan Universal PCR Master Mix (Applied Biosystems). The sample mix and TaqMan assays were loaded separately into the wells of 96.96 Gene expression Dynamic Arrays (Fluidigm) in presence of appropriate loading reagents. The arrays were read in a Biomark analysis system (Fluidigm). ΔCt values were calculated in reference to the average of *Atp5a1, Hprt1* and *Ubc*.

The expression of each gene was fit by a bivariate normal distribution (R mixtools package (*58*)) and a cut-off set at the average position between the lower and upper gaussian midpoints. Genes above this were considered expressed if above this cut-off. qPCR levels and metadata are provided in Supplementary File S1.

#### *S. mansoni* C1 single cell RNA-seq and analysis

For *S. mansoni*, three small (5–10 μm) C1 Single-Cell Auto Prep IFC chips (Fluidigm) were primed and 5000 cells were sorted directly into the chip. To allow estimation of technical variability, 1 μl of a 1:4000 dilution of ERCC (External RNA Controls Consortium) spike-in mix (Ambion, Life Technologies) was added to the lysis reagent. Cell capture sites were visually inspected one by one using a microscope. The capture sites that did not contain single cells were noted and were removed from downstream analysis. Reverse transcription and cDNA preamplification were performed using the SMARTer Ultra Low RNA kit (Clontech) and the Advantage 2 PCR kit according to the manufacturer’s instructions on the C1 device. cDNA was harvested and diluted to 0.1–0.3 ng/μl and libraries were prepared in 96-well plates using a Nextera XT DNA Sample Preparation kit (Illumina) according to the protocol supplied by Fluidigm. Libraries were pooled and sequenced on an Illumina HiSeq2500 using paired-end 75-bp reads.

Salmon was used to estimate gene expression counts (*59*). Poor quality libraries were eliminated using Scater (*60*) based on exonic and mitochondrial read counts. For all queries, a gene was considered expressed if the Log10 (1+normalized count) was above 0.5. Gene overlap was tested using Fisher’s method. Counts and condition matrices are provided in Supplementary File S3.

#### *N. brasiliensis* C1 single cell RNA-seq

T cells from *N. brasiliensis* infected mice were captured with C1 analogously to the *S. mansoni* mice. The cells were sorted for CD3+CD4+SELL-, from mesLN, medLN, lung and spleens. This data is available but was not used due to low quality.

#### *N. brasiliensis* Smart-seq2 single-cell RNA-seq and analysis

For this analysis we included CD3+CD4+SELL-cells from the lungs at 30 days after infection, CD3+CD4+Rora^teal^+ and CD3+CD4+Rora^teal^-cells from *N. brasiliensis* infected *Rora*^*+/teal*^ reporter mice. Further we include cells from lungs, spleens, medLN and mesLN from infected and uninfected *Rora*^*+/teal*^ mice, 7 days after infection.

Single cell transcriptomes were generated by using the Smartseq-2 protocol (*61*). Single cells were sorted into 96 well plate that contained 5μl of of Triton-X lysis buffer, 1 µl of 10 µM olido-dT30-VN, 1 µl dNTP mix (25 mM each) and ERCC controls at a final dilution of 1:64m and immediately stored in -80°C. cDNA was submitted for the amplification for 25 cycles on an Alpha Cycler 4 thermal cycler. Amplified cDNA went through two subsequent rounds of cleaning by using Agencourt AMPure XP beads (Beckman Coulter UK Ltd, High Wycombe, UK) at a 1.0x ratio on a liquid handler Zephyr G3 NGS Workstation (PerkinElmer) and eluted in 20ul of RNAse free water. Purified cDNA was subjected to quality control using 1 µl of cDNA on an Agilent 2100 BioAnalyser (Agilent Technologies, Santa Clara, CA, USA) using the Agilent High Sensitivity DNA kit. Samples were normalised to a concentration of 0.3 ng/µl. Nextera libraries were prepared using Nextera XT DNA Sample Preparation kit (Illumina) according to the protocol supplied by Fluidigm. The library preparation has been done in combination with benchtop liquid handlers: Zephyr G3 NGS Workstation (PerkinElmer) and Mantis (Formulatrix). Purified pool of Nextera libraries was subjected to quality control using 1 µl of cDNA on an Agilent 2100 BioAnalyser using the Agilent High Sensitivity DNA kit.

Salmon v0.8.2 (*59*) was used to map the reads to ensembl cDNA reference GRCm38 to quantify the gene expression. Counts were used for all the analyses. Low quality cells with >15% mitochondrial genes or <1500 total genes were filtered out. Seurat (*62*) was used for single-cell analysis: Highly variable genes were calculated (x.low.cutoff=0, x.high.cutoff=5, y.cutoff=0.1), PCA performed on these (pcs.compute=50), and t-SNE on the first 25 PCA components. Finally clustering was done (reduction.type=“pca”, dims.use=1:25, resolution=0.5). Common marker genes were plotted on top of the clusters and the cluster identities were assigned manually (*Gata3, Tbx21, Foxp3, Sell*, and *Rorc*). The TCR sequences for each single T cell were assembled using TraCeR (*63*) which allowed the reconstruction of the TCRs from scRNA-seq data and their expression abundance, in addition to identification of the size, diversity and lineage relation of clonal subpopulations. Cells for which more than two alpha or beta chains were identified were excluded from further analysis. iNKT cells were detected by their characteristic TCRA gene segments (TRAV11–TRAJ18). Only CD4+ T cells were kept in the final plot; other cells remain annotated in the ArrayExpress submission. Counts and condition matrices are provided in Supplementary File S2.

### Screening for upstream regulators of *Rora*

#### Cell culture

Naive CD4+ T helper cells were extracted from spleens of JaxJ mice, 8 weeks old males (Stem Cell Technologies #19765), according to the manufacturer’s protocol. Up to 4 spleens were pooled in each biological replicate. The cells were activated in 96-well round bottom plates that were coated with 3ul/ml anti-CD3e (BioLegend #100202) and 5ul/ml anti-CD28 (BioLegend #102102) for 4 hours at 32°C. 24 hours later cytokines were added (See Supplementary file S4). 72 hours after activation the cells were pelleted, resuspended in RLT buffer (Qiagen #79216), and stored at -80°C.

For the repeated screen, culturing was done in the same manner. The cells were then stained with PI and anti-CD4 eFluor 660 (eBioscience 50-0041-82, clone GK1.5) for 30 minutes in IMDM media. The cells were then FACS sorted to retain live CD4+ T cells, on average 5000 cells per well.

During Th0 conditions, it is possible for the cells to become somewhat Th1 polarized; we have looked at markers in the data but can neither confirm nor disprove that this is the case for these experiments.

#### RNA-sequencing

For the first screen, RNA was extracted using Agencourt AMPure XP beads (Catalog #A63881) at 1.5:1 ratio into 10ul elution volume. RNA-seq was done following Smart-seq2 (*61*) but with the following changes: The input was increased to 7ul and Mix 1 consisted of 2ul 10uM dNTP and 2.5ul 10uM oligo-dT. Mix 2 consisted of 5ul SmartScribe buffer, 5ul 5M Betaine, 1.25ul SmartScribe, 1.25ul (100uM) DTT, 0.15ul 1M MgCl2, 0.38ul 100uM TSO, and 0.47ul SUPERase inhibitor. SPRI cleanup was done at ratio 0.8:1 following first-strand synthesis, eluted into 12ul NFW. Mix 3 consisted of 12.5ul KAPA HiFi 2x master mix and 0.5ul 10uM ISPCR. PCR was done with 10 cycles. In the second screen, RNA-seq was done similarly with the following modification: No SPRI-cleanup was done after first-strand synthesis. Instead a larger amount of mix 3 was added directly.

On average 0.2ng cDNA was then used for Nextera XT library preparation, done at 25% of volume recommended by manufacturers protocol. Sequencing was done with 2 lanes of Illumina 50bp PE HiSeq 2500.

#### Upstream analysis

Salmon was used to estimate gene expression counts. Poor quality libraries were eliminated using Scater based on exonic and mitochondrial read counts. Genes with average normalized counts below 0.3 were removed. The increase/decrease in *Rora* expression level was computed by DEseq2 (*52*) using a linear model: normalized counts ∼ treatment + mouseReplicate, where treatment is a factorial variable. The correlation heatmap between treatments was calculated as cor(cor(FC)). The first correlation is used as a dimensionality reduction while the second correlation is used to remove the technical bias toward positive correlation. Clustering was done using R hclust, standard parameters.

The CellTrace FACS data was analyzed with FACSanadu (*64*) to obtain cells in each division stage. The cell counts were analyzed across replicates and compared to controls using R.

### Metabolite LC-MS

Metabolites were extracted using a modification of a previous method (*65*). Briefly, roughly 200k cells were resuspended in chloroform/methanol (2:1). After mixing samples, water was added (300 μL) and centrifuged at 13,000 x *g* for 20 min. The organic layer (lower fraction) was collected and dried under a stream of nitrogen. The organic fraction was reconstituted in chloroform/methanol (1:1; 100 μL) and 10 μL were mixed in 90 μL IPA/acetonitrile/water (2:1:1). The samples were analysed by LC-MS using an LTQ Orbitrap Elite Mass Spectrometer (Thermo Scientific). Sample (5 μL) was injected onto a C18 packed-tip column (75μm x 100mm, Thermo Scientific) at 55°C. The mobile phase A was acetonitrile/water 60:40, 10 mM ammonium formate whilst B was LC–MS-grade acetonitrile/isopropanol 10:90, 10 mmol/L ammonium formate. In negative ion mode, ammonium acetate was used to aid ionisation. In both positive and negative ion mode, the gradient used (Supplementary File S4) was run at a flow rate 0.5 mL/min. Data were acquired with a mass range of 100-2000 m/z. Chromatograms were converted to mzML format and annotated using an in-house R script. The annotated metabolites were size factor normalized to enable comparison. PCA was performed on the *in vitro* and *in vivo* samples separately (Figure S5). Lungs and spleens from *in vivo* samples appear to cluster together, suggesting no need to correct for tissue differences. *In vitro* samples were compared with Limma (*66*), with all naive cells compared to all day 6 samples Th0/1/2. For the *in vivo* samples, we used the simplest model of ∼1+isko. Bayesian error correction was performed.

## Supplementary Figures

**Supplementary Figure 1:**
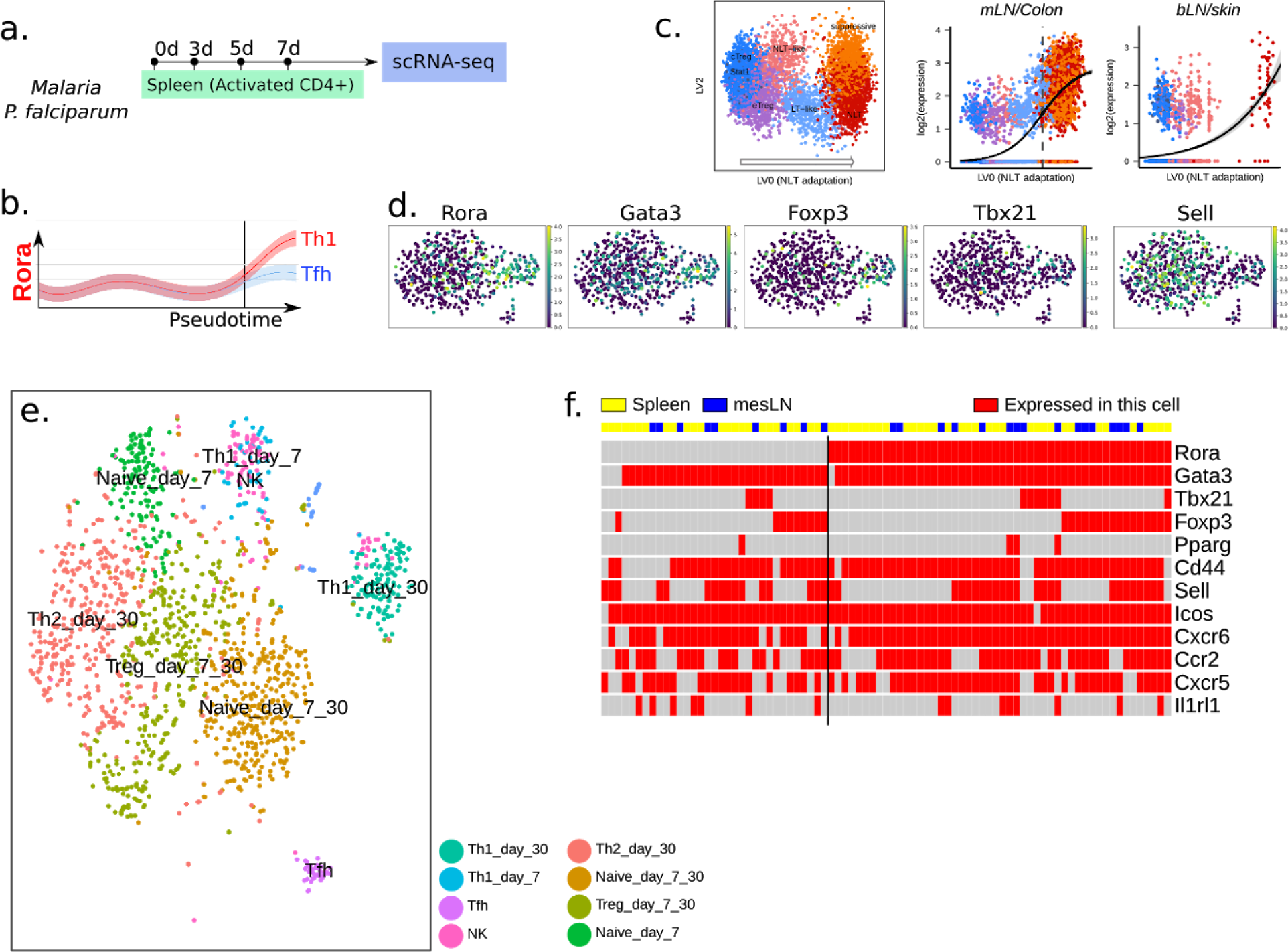
**(a)** Experimental design for a previous malaria *in vivo* time course scRNA-seq experiment (*1*), and **(b)** pseudo-time reconstruction of *Rora* gene expression, in Th1 and Tfh. **(c)** Panels reproduced from a study comparing Treg cells from lymphoid and non-lymphoid tissue (*2*). This study shows that Rora is expressed mainly in peripheral-like (IL10-producing) Tregs, and non-lymphoid tissue Tregs. **(d)** Expression of *Rora*, and other marker genes, in CD4 T cells in mouse melanoma (*3*). This dataset is in agreement with our conclusion that *Rora* is expressed mainly in activated T cells. **(e)** Clustering of all cells related to the *N. Brasiliensis* scRNA-seq experiment. **(e)** Expression level (single cell RNA-seq) of selected genes in activated CD62L-/CD44+ T cells, 6 weeks after an *S. mansoni* infection, confirming *Rora* presence. Note that because activated CD4+ T cells were sorted for this experiment, the overlap with Rora+ and Sell-should be less than in the previous *N. brasiliensis* experiment.

**Supplementary Figure 2:**
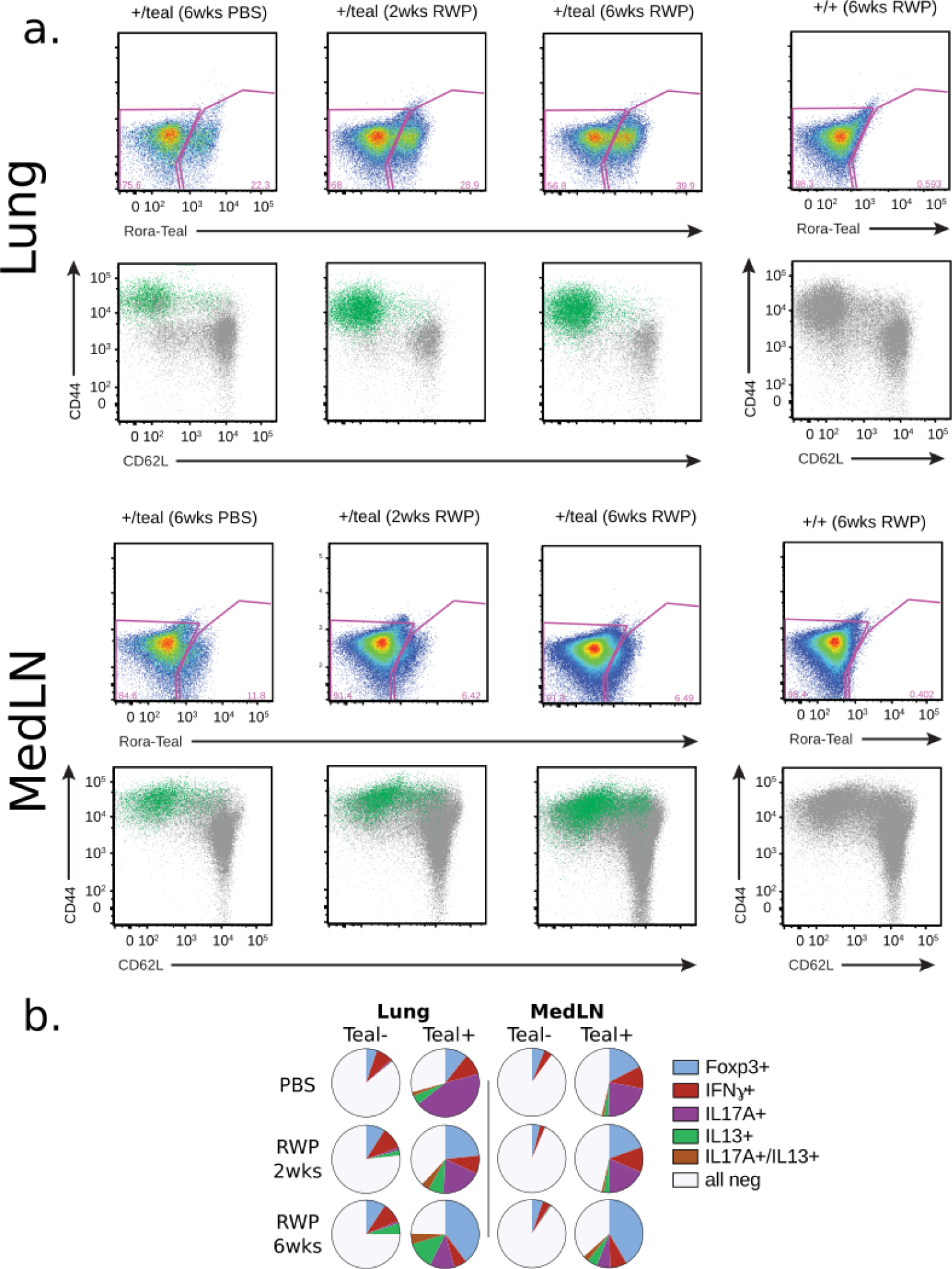
**(a)** FACS analysis of Rora+ and Rora-CD4+ T cells from RWP treated mice. **(b)** Analogous plot to Figure 3c, except IL17 and IFN are also included.

**Supplementary Figure 3.**
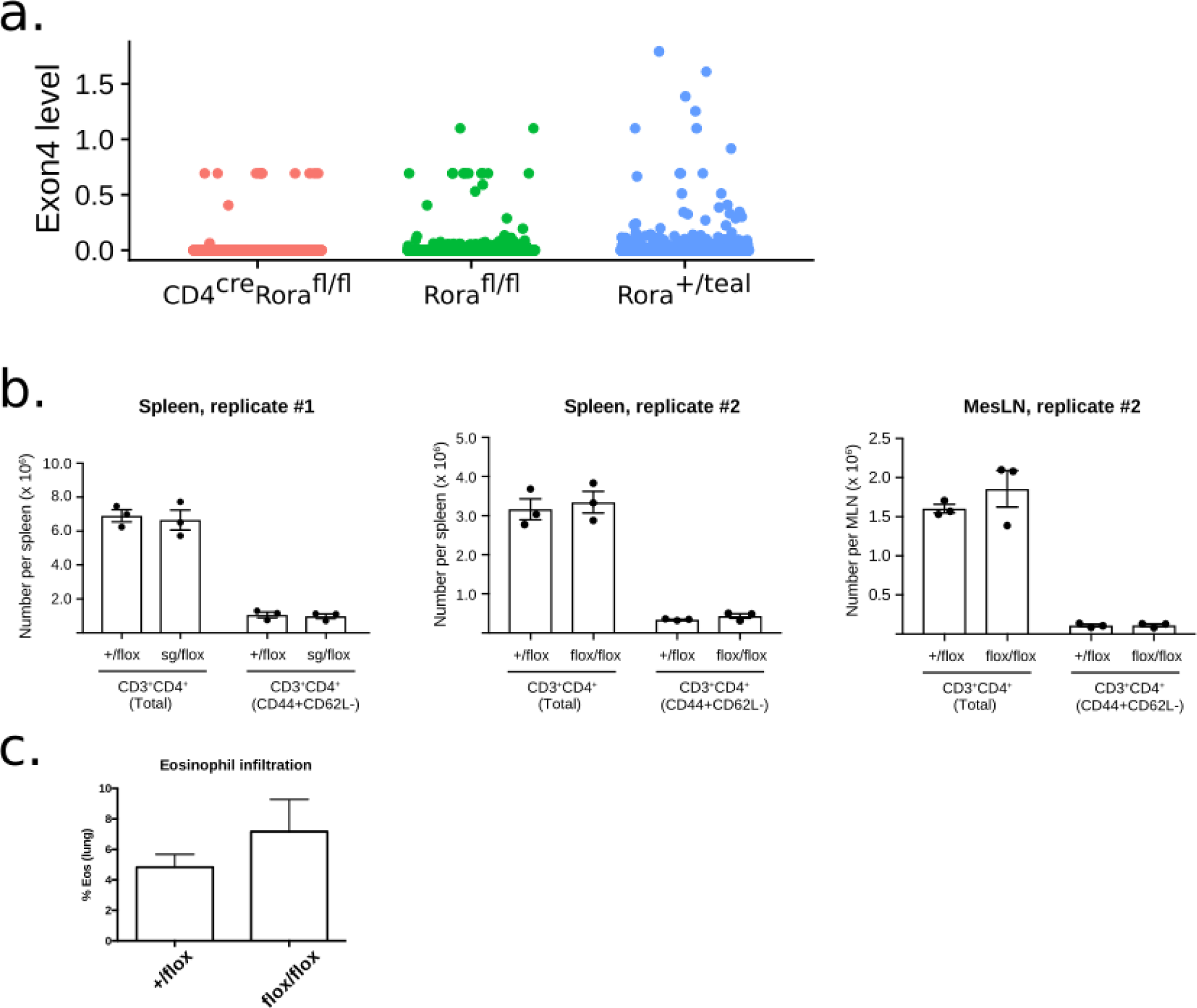
**(a):** Usage of exon 4 in *Rora* KO (red) *vs* control mouse (green and blue), based on scRNA-seq data. The relative exon usage is calculated as Log[(exon4_reads+1)/(rora_reads+1)]. Exon 4 is conditionally removed in the KO mouse. Note that due to intrinsic noise in scRNA-seq data, the presence of exon 4 cannot be assessed for many of the cells. **(b)** CD4+ T cell count does appear affected in Cd4^Cre^*Rora*^*fl/fl*^ KO vs control under non-infected conditions. **(c)** Eosinophil infiltration level in KO vs control, as indicated by a manual scoring.

**Supplementary figure 4:**
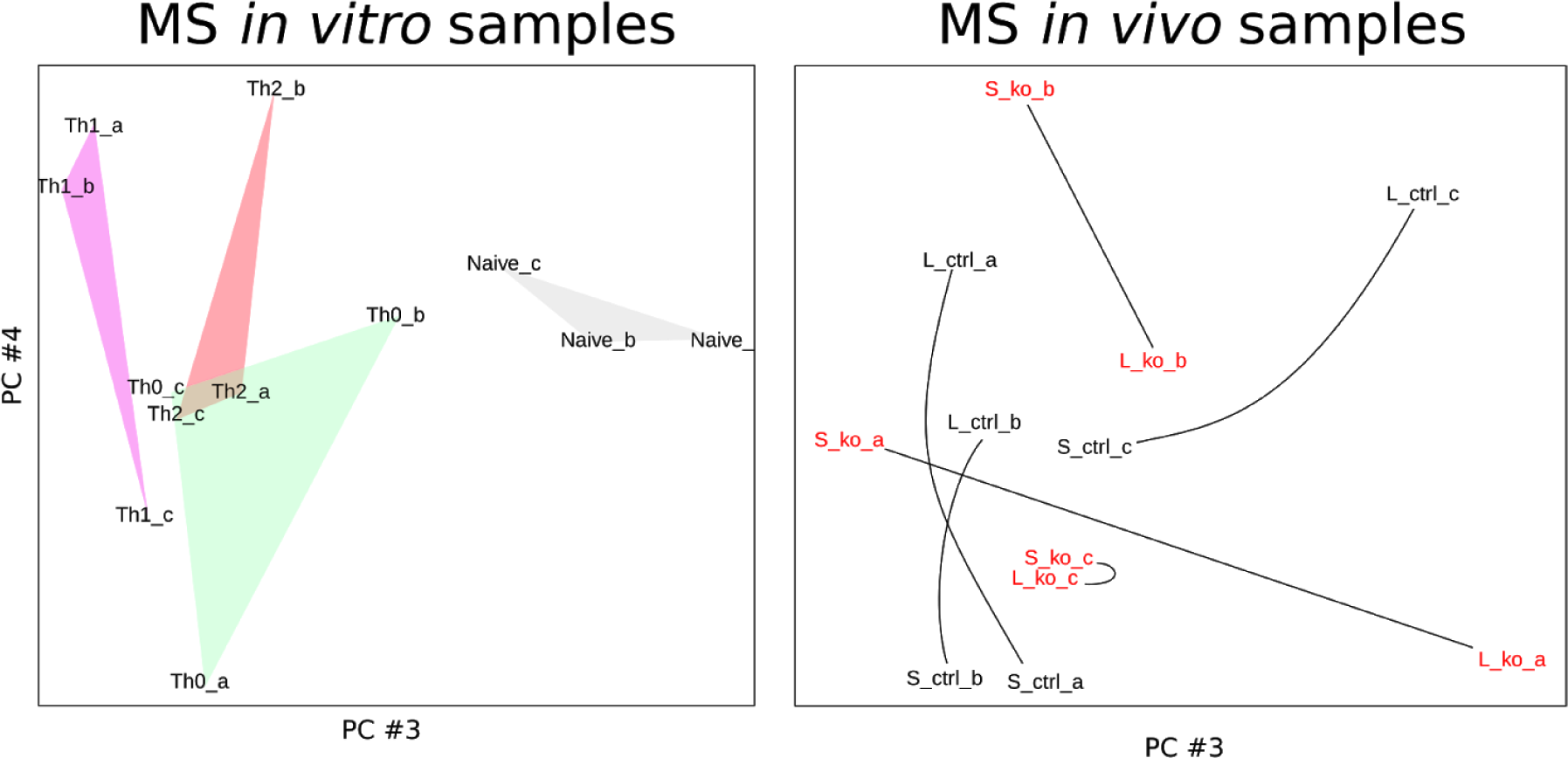
PCA analysis of the LC-MS samples. *In vitro* samples separate into types. *In vivo* samples appear to primarily cluster on donors, with spleen and lung from mouse ko_c being the most clear example.

**Supplementary figure 5:**
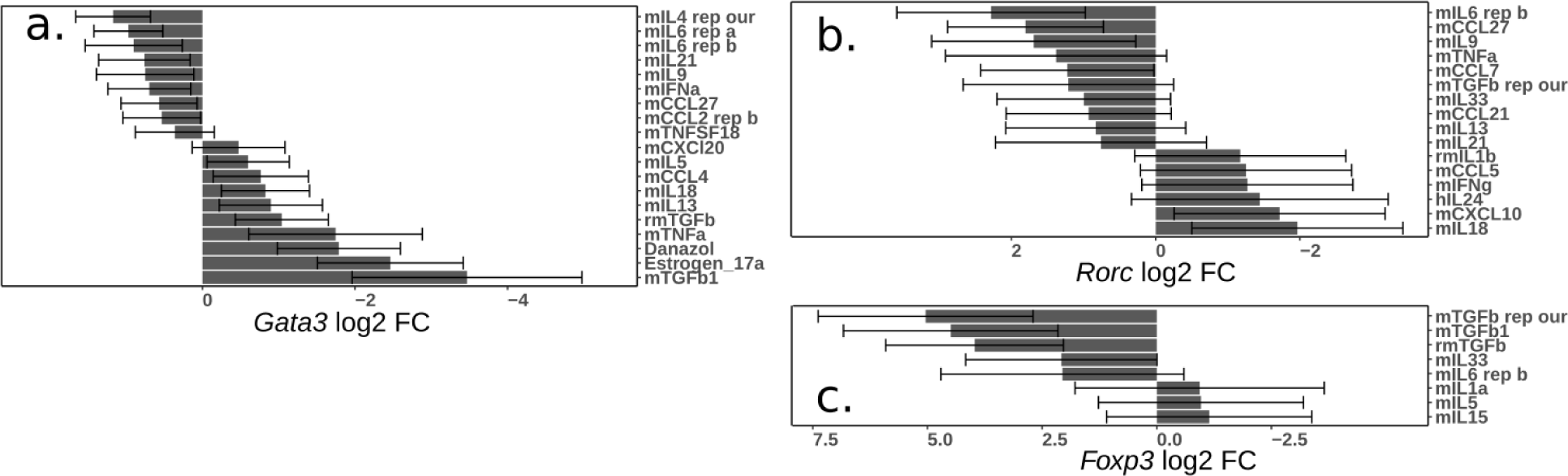
Quality control of the database of cytokine-treated CD4+ T helper cells. **(a)** Regulators of Th2 cells. **(b)** Regulators of Th17 cells. **(c)** Regulators of Treg cells.

