## Supplemental File 2 for "*Rora* regulates activated T helper cells during inflammation": explanation_rnaseq.txt

The data structure here follows largely from the way we submitted samples for sequencing, as well as how the provided code operates on it. This led to a somewhat convoluted organization - please see the description files in subfolders to see how the condition matrices should be interpreted
